# Breaking the culture habit: metagenomic diagnosis of companion animal skin infections

**DOI:** 10.1101/2025.09.26.678841

**Authors:** Andrea Ottesen, Brandon Kocurek, Mark Mammel, Sanchez St. Laurent, Jaclyn Dietrich, Sarah Pauley, Stephen Cole, Shelley Rankin, Olgica Ceric

## Abstract

**Background:** Skin infections have been described as the primary cause for presentation in veterinary small animal practices and they frequently result in prescription of both topical and systemic antibiotics. Because such infections are often secondary complications of other underlying pathologies, recurrent infections are common and can lead to multiple antibiotic exposures. This scenario creates steady selection pressure toward antibiotic resistance at the confluence of the skin (the largest mammalian organ), the bloodstream, and shared human and animal environments. This case study compares metagenomic (MGX) data with aerobic culture to evaluate diagnostic utility for simultaneous identification and characterization of pathogens, microbiomes, and resistomes of companion animal skin infections.

**Results:** One feline and eight canine skin swabs were analyzed with aerobic culture and traditional antimicrobial susceptibility testing (AST) and compared with MGX profiling. Veterinary laboratory diagnostic (VDL) culture and AST identified *Staphylococcus aureus, S. pseudintermedius*, *S. schleiferi,* methicillin resistant (MR) *S. schleiferi* (MRSS), MR *S. pseudintermedius* (MRSP) and *Pseudomonas aeruginosa* from skin swabs. MGX data described the identical bacterial pathogens recovered by aerobic culture and methicillin resistance genes *mecA, mecI, mecR1* in samples for which AST confirmed MRSP and MRSS. MGX data also identified *mec* genes in samples without culture-based confirmation of MR phenotypes. MGX data also described multi-domain composition of microbiomes of infected skin including bacteria, fungi, viruses, phages, AMR, plasmids, and metabolic features associated with skin infections.

**Conclusions:** MGX data identified the identical pathogens and inferred AMR phenotypes as culture-based diagnostic testing, and additionally characterizedo multi-domain microbiota, mobile AMR elements, and metabolic features. Efforts to accelerate cures by precision medical responses depend on accelerated precision diagnostics. Challenges remain for the implementation of MGX data into veterinary diagnostic laboratory investigation and response. We demonstrate with a small case study, that MGX data can be used to complement current state of the art VDL results and potentially advance a judicious veterinary medical response regarding antibiotic administration for companion animal skin infections. In the future, simultaneous description of the polymicrobial ecology of skin infections (bacterial, viruses, phages, fungi, and even functional metabolomic features) provided by MGX data can advance epidemiology, develop new treatment strategies, accelerate diagnostics and provide data for artificial intelligence (AI) models focused on advancing veterinary diagnostics and medical treatments.

## Introduction

Companion animals’ skin infections have been described as the primary reason for presentation in small animal practices and result in frequent antibiotic administration(1). Skin infections are often secondary complications of reduced immunity associated with underlying pathology and/or dysbiosis and precise diagnoses can be challenging. It is not uncommon for skin infections to reoccur, and this can result in subsequent re-exposure to antibiotics. This scenario results in AMR selection pressures at the intersect of skin (the largest mammalian organ), the bloodstream, and shared human and animal environments(1). Additionally, dogs and cats are often treated with the same antibiotics used in human medicine such as fluoroquinolones, penicillins, and cephalosporins(2). This, considered with the proximity shared by humans and companion animals underscores the importance of AMR stewardship at this important interface (3). To reduce selection pressures for Difficult to Treat Resistance (DTR)^1^ pathogens, diagnostics that identify pathogen, resistome, and total polymicrobial ecology in a single pass, will be transformatory to public health(4).

MGX data have been used to identify pathogens associated with etiologies of unknown origin in human and veterinary medicine (5–9) for more than two decades. Numerous labs have integrated MGX options into plant, animal, and human pathology diagnostics due to the sheer practicality of results that describe the complete landscape of an infection microbiome (infectome). Diagnostics for viral pathogens were some of the first to implement MGX methods, due to challenges associated with culturing viruses(8). Kaszab et al. describe a shift from targeted molecular assays to MGX diagnostics for identification of viruses causing illness and economic loss in the farm animal industry(10). MGX data can also be used for genome focused characterization as illustrated in a retrospective investigation of Tyzzer disease in foals, which described pathovar and virulence of *Clostridium piliforme* from shotgun sequence data from deceased animals(11).

All assays are most powerful when optimized for a specific objective, which is no different for MGX-based diagnostics. This is especially significant for the veterinary diagnostics where multiple host species and their associated microbiomes, resistomes, and pathologies, provide a uniquely complex challenge for medical practice. Many researchers have been advancing the field with MGX applications for detection of porcine virus(12, 13), bovine respiratory disease (14), periweaning failure-to-thrive syndrome(15), shaking mink syndrome(8), meningoencephalomyelitis of unknown origin(16), cat scratch disease(17), neuroleptospirosis(9), and many more. In this study, we demonstrate with a small case study how MGX data can be used to advance precision diagnostics and a judicious treatment response as well as pathogen epidemiology and zoonotic risk assessment.

## Results

### Veterinary diagnostic laboratory (VDL) culture results

VDL aerobic culture results from swabs of skin infections of one cat and eight dogs identified *Staphylococcus schleiferi, Staphylococcus aureus,* methicillin resistant *Staphylococcus pseudintermedius* (MRSP), methicillin resistant *Staphylococcus schleiferi* (MRSS) and *Pseudomonas aeruginosa* (Table 1).

**Table 1.** VDL culture and AST results for skin infections from 1 cat and 8 dogs.

| ID # | Organism Recovered | Resistance to Antibiotic (I) Intermediate Resistance |
| --- | --- | --- |
| FS1 | *Staphylococcus aureus* | Cefovecin (I), Cefpodoxime (I), Enrofloxacin |
| CS1 | *Pseudomonas aeruginosa* | Cefovecin, Ceftiofur, Ciprofloxacin (I), Enrofloxacin (I), |
| CS1 | *Staphylococcus schleiferi* | Enrofloxacin |
| CS2 | *Pseudomonas aeruginosa* | Cefovecin, Ceftiofur, Ciprofloxacin (I), Enrofloxacin (I), |
| CS3 | *Pseudomonas aeruginosa* | Cefovecin, Ceftiofur, Ciprofloxacin (I), Enrofloxacin (I), |
| CS4 | *MR Staphylococcus schleiferi* | Cefovecin, Clindamycin, Cefpodoxime, Doxycycline, Erythromycin, Enrofloxacin (I), Marbofloxacin (I), Oxacillin, Benzylpenicillin |
| CS5 | *MR Staphylococcus pseudintermedius* | Doxycycline, Enrofloxacin, Gentamicin (I), Minocycline, Marbofloxacin, Oxacillin, Benzylpenicillin, Pradofloxacin, Trimethoprim/Sulfamethoxazole |
| CS6 | *Staphylococcus pseudintermedius* | Doxycycline, Minocycline, Benzylpenicillin |
| CS7 | *Staphylococcus pseudintermedius* | Benzylpenicillin |
| CS8 | *MR Staphylococcus schleiferi* | Cefovecin, Cefpodoxime (I), Enrofloxacin, Marbofloxacin, Oxacillin, Benzylpenicillin, Pradofloxacin (I) |

Full antibiotic sensitivity testing profiles available in Supplementary Materials S1.

### MGX profiling

MGX data identified identical pathogens to those recovered by culture methods (Figure 1B) while simultaneously describing multi-domain microbiota (bacteria (Figure 1C), virus, phage (Figure 2), fungi (Figure 3), protist, metabolic features, and resistomes (antimicrobial resistance genes (ARGs) and plasmids for each sample (Figures 5-8). Culture for samples CS4 and CS5 identified methicillin resistant *Staphylococcus schleiferi* (MRSS) and methicillin resistant *Staphylococcus pseudintermedius* (MRSP) respectively. MGX data identified the same bacterial species (*S. schleiferi* and *S. pseudintermidius*) and methicillin resistance gene determinants *mec*A in sample CS4, and *mec*A, *mec*R1, and *mec*I in CS5. Although not reported by the VDL, MGX and quasimetagenomic data (qMGX)(18–20) suggest that FS1 and CS1 and potentially other samples may have also been positive for methicillin resistant species of *Staphylococcus* that were not reported by the VDL aerobic culture assays.

**Figure 1. Culture and AST results with MGX characterized pathogens, AMR, and polymicrobial composition of skin infections for 1 feline and 8 canine skin samples**

**Figure 1.**
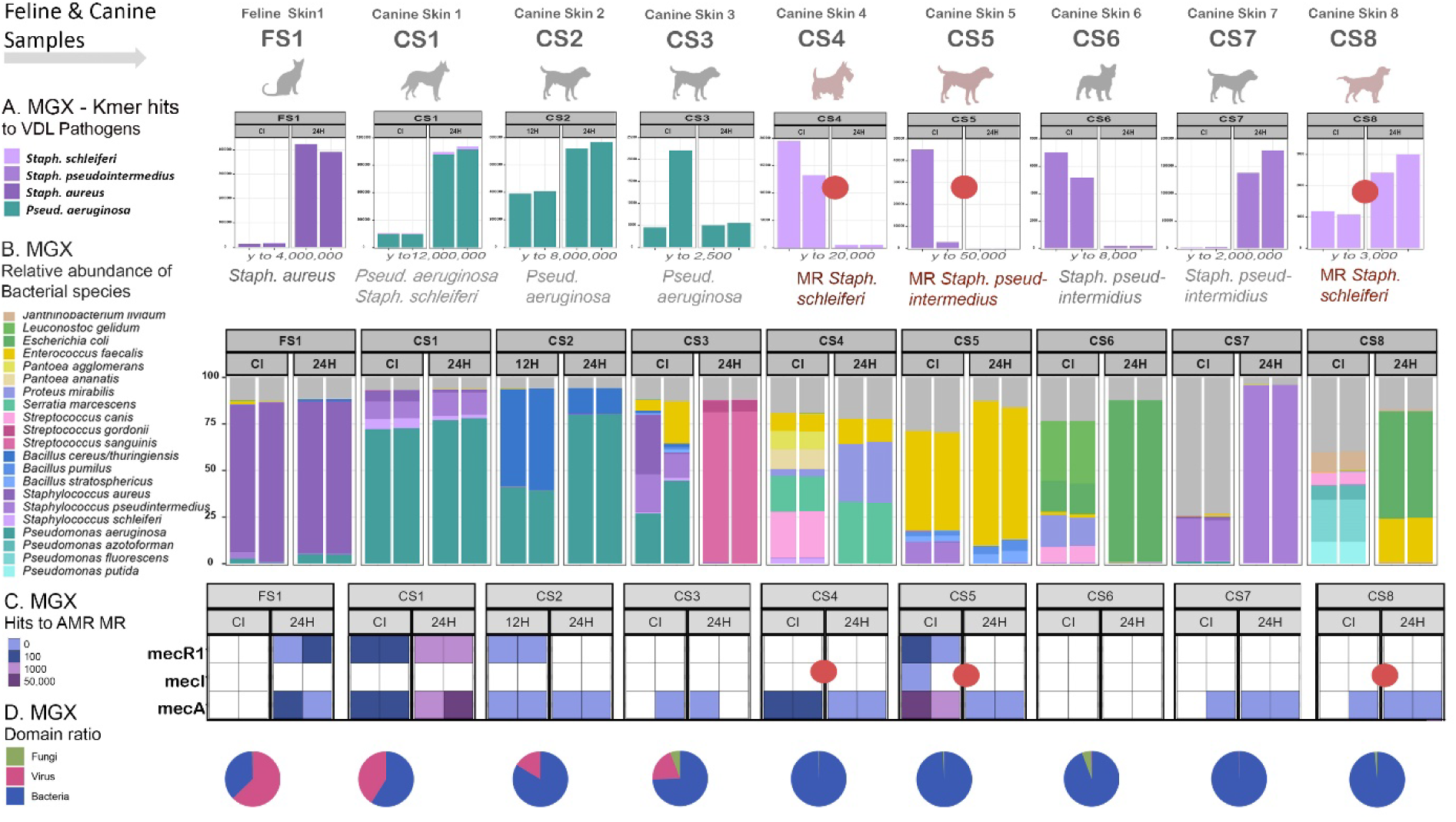
VDL Aerobic Culture & AST, MGX-pathogen detection, MGX-AMR MR, identification and MGX polymicrobial composition, in 1 feline and 8 canine skin samples. **A.** VDL aerobic culture & AST pathogen recovery results reported methicillin (MR) resistant phenotypes from feline and canine skin infections (CS4, CS5, &CS8) are shown with MGX hits to the same species. **B.** MGX and qMGX relative abundance of all bacteria identified in companion animal skin infections for independent replicates of culture independent MGX samples (CI) and qMGX (shotgun sequencing of enrichments) at specific temporal timepoints Ie; (12, 24). *For sample CS2, insufficient DNA from CI was recovered and only 12H qMGX data could be used for reporting. **C.** MGX AMR – Methicillin resistance determinants: *mec*A, *mec*R1, and *mec*I **D.** MGX – Multi-domain ratio for viruses, fungi, and bacteria observed across all annotated reads.

MGX descriptions of bacterial microbial community composition are presented in relative abundance (Figure 1B), which considers number of reads per sample and provides ‘normalized’ reports of taxonomic composition. Hits to pathogens are presented as actual ‘hit’ to the species (Figure 1A). Figure 1 shows 2 replicates of metagenomic data and 2 replicates of quasimetagenomic data or enriched (Universal Pre-enrichment Broth (UPB)) samples. In certain pathogen and AMR detection efforts, enriched shotgun data provide more information than metagenomic data alone(18–21).

### Staphylococcus and Pseudomonas in skin infections

Previous culture based veterinary diagnostics have described *Staphylococcus pseudintermedius* as the most frequently recovered Gram-positive species from canine skin infections (65%) (n = 171)(22). This ratio aligned with results observed here. VDL aerobic culture to identify pathogens of known clinical significance such as *Staphylococcus spp.* may overlook co-occurring taxa which may or may not play a role in infections. In the small number of canine skin infections examined here (n=8), *S. pseudintermedius* was recovered from three samples (CS5, CS6, and CS7). In samples from which *S. pseudintermedius* was recovered, predominant Gram-positive members of the community included *Enterococcus faecalis* (CS5), *Leuconostoc gelidum,* and *Streptococcus canis* (CS6). *Staphylococcus schleiferi* was recovered by VDL testing from three canine infections but represented a low abundant member of the community by MGX profiling. Several *Staphylococcus* spp. co-occurred in infections including *S. aureus, S. pseudintermedius, S*. *schleiferi* and *S. saprophyticus*. Additional *Staphylococcus* species observed in low abundance included *S. capitis, S. epidermidis, S. hyicus, S. pasteuri, S. simulans, S. succinus, S. virulinus, S. warneri,* and *S. xylosus*.

*Pseudomonas aeruginosa* has also been previously described as the most frequently recovered Gram-negative species(22) from canine skin infections. This observation aligned with feline and canine samples examined here from which *Pseudomonas aeruginosa* was also the most abundant taxon in MGX infection profiles. Other dominant Gram-negative species by sample included *Serratia marcescens, Pantoea ananatis, Pantoea agglomerans* (CS4), and *Escherichia coli* (CS6). MGX data also described numerous additional *Pseudomonas* species that may or may not play a role in skin infections. In total, hits to more than 100 *Pseudomonas* species were described. The top ten, ranked by abundance included *Pseudomonas aeruginosa*, *P. azotoformans, P. fluorescens*, *P. rhodesiae, P. coleopterorum, P.putida, P. protegens, P. proteolytica, P. trivialis,* and *P. extremorientalis* (Table 2).

**Table 2.** Ten most abundant *Pseudomonas* species by unique k-mer hits in MGX data of skin samples.

| *Pseudomonas* species | FS1 | CS1 | CS2 | CS3 | CS4 | CS5 | CS6 | CS7 | CS8 |
| --- | --- | --- | --- | --- | --- | --- | --- | --- | --- |
| *Pseudomonas aeruginosa* | 4771 | 1443264 | 3992762 | 13285 | 4908 | 10290 | 2864 | 6350 | 3272 |
| *Pseudomonas azotoformans* | 1 | 3 | 108 | 0 | 50 | 266 | 3 | 1 | 154989 |
| *Pseudomonas fluorescens* | 1 | 12 | 151 | 0 | 927 | 148 | 6 | 184 | 148889 |
| *Pseudomonas rhodesiae* | 0 | 2 | 82 | 0 | 65 | 1 | 3 | 0 | 121802 |
| *Pseudomonas coleopterorum* | 10 | 3 | 15 | 0 | 62550 | 9 | 4 | 0 | 15897 |
| *Pseudomonas putida* | 1 | 14 | 127 | 0 | 366 | 21 | 2 | 178 | 67817 |
| *Pseudomonas protegens* | 0 | 2 | 39 | 0 | 54 | 3 | 0 | 0 | 52601 |
| *Pseudomonas proteolytica* | 0 | 0 | 14 | 0 | 29 | 15 | 1 | 0 | 20634 |
| *Pseudomonas trivialis* | 0 | 1 | 20 | 0 | 140 | 8 | 1 | 2 | 15129 |
| *Pseudomonas extremorientalis* | 0 | 1 | 89 | 0 | 37 | 1 | 0 | 1 | 10641 |

### MGX: Viruses and Phages

MGX data provided robust DNA virus and phage characterization in addition to pathogen detection/identification and polymicrobial microbiome profiling. Viruses and phages observed in feline and canine skin infections are described in Figure 2. The most dominant phage observed was *Pseudomonas* phage Pf1, which correlates with prevalence of *Pseudomonas* across the samples. Torque teno canis virus was observed in Canine skin infection sample CS6.

**Figure 2.**
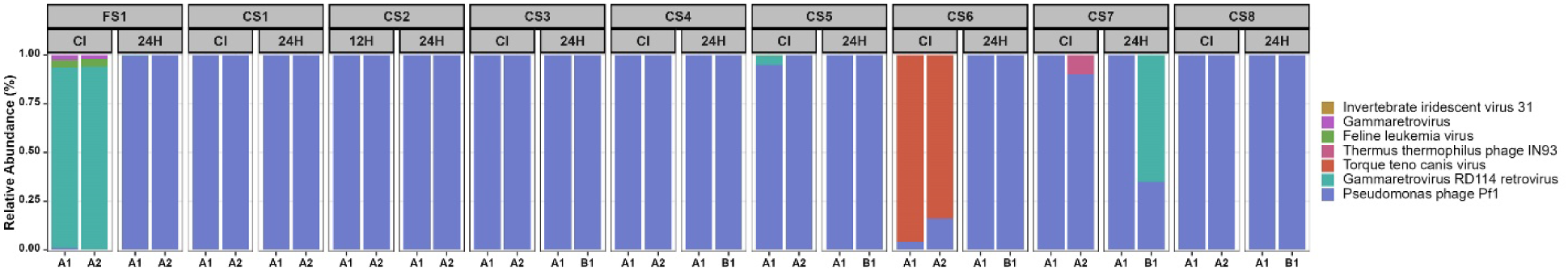
MGX-Profiles of Viral and Phage species observed in feline and canine skin infections.

### MGX-Fungal Taxa observed in MGX data from feline and canine skin samples

*Alternaria*, *Malassezia, and Aspergillus* have been described as the most commonly observed fungal genera associated with allergy and infection in animals (23) although we only observed *Malassezia* and a very low abundance of *Aspergillus* in the small number of samples examined here. *Malasseszia pachydermatis* was observed in samples CS1 and CS8 (very low abundance in CS8) (Figure X). Other fungal taxa observed in MGX data included *Mucor racemosus* (24), *M. plumbeus*, *Pseudozyma thailandica, Melampsora, Balansia obtecta, Puccinia, Plasmopara, Neonectria, Astraeus odoratus, and Tubaria*.

**Figure 3.**
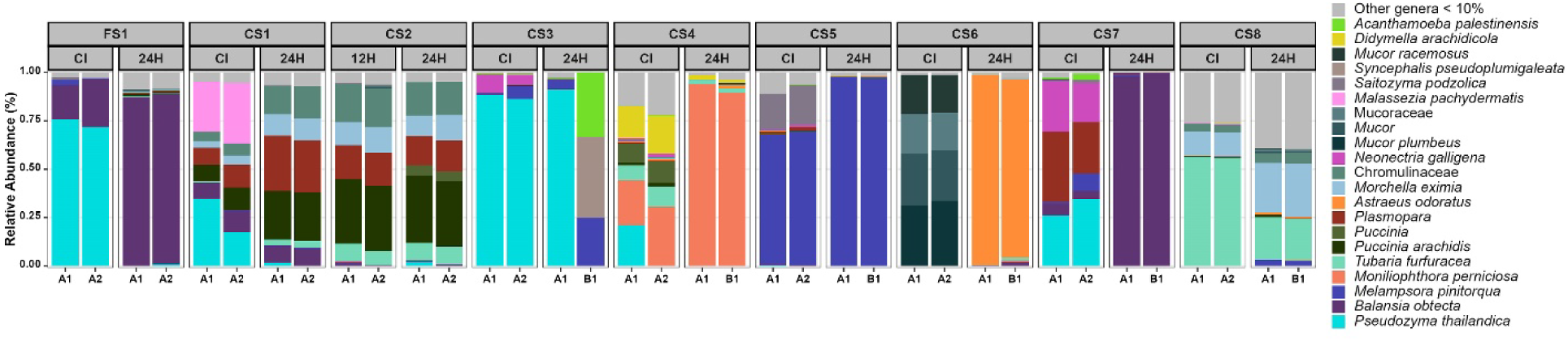
MGX-Fungal Taxa observed in MGX data from feline and canine skin samples.

### MGX-Resistome

Multiple classes of critically important antimicrobial resistance (25) were identified by MGX data. Those included macrolide resistance genes *acr*B, *erm*B, *ermT*, *erm*A, *mpb*H, and beta-lactam genes: CTX-M-104, CTX-M-14, CTX-M-27, CTX-M-65, and CTX-M-9, TEM 207 Aminoglycosides, fluoroquinolones and macrolides, lincosamides and streptogramins. Panels X-X show the number of hits to various ARGs across the feline and canine skin samples.

**Figure 4.**
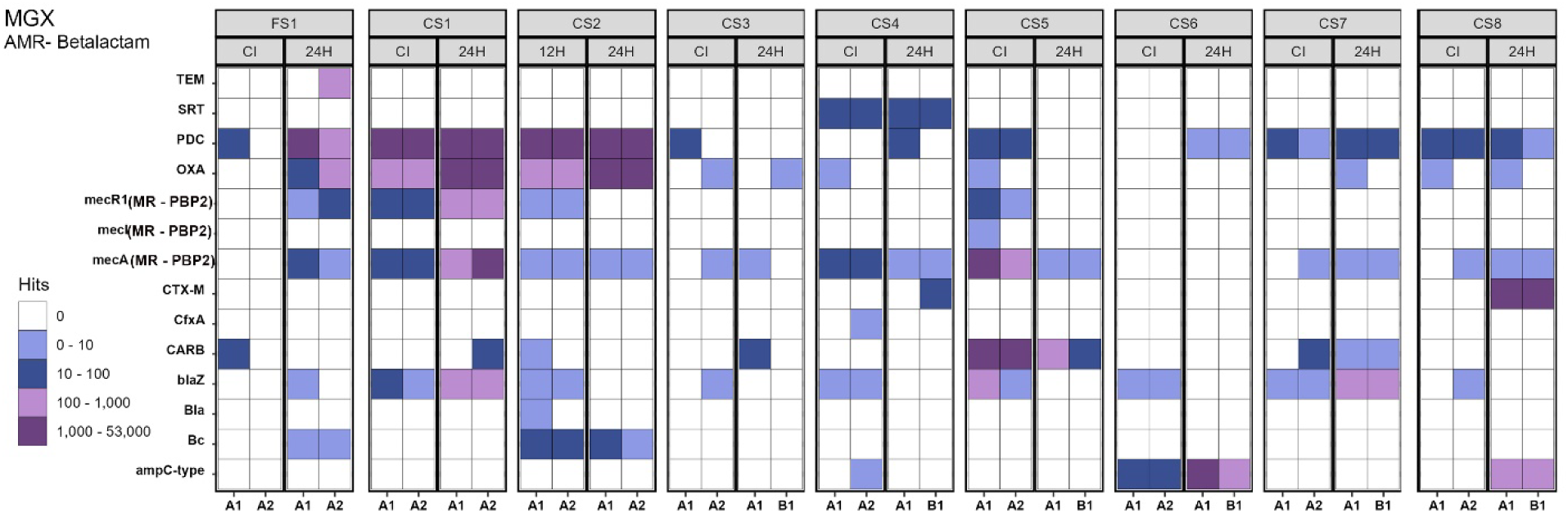
MGX Betalactam ARGs in feline and canine skin samples.

**Figure 5.**
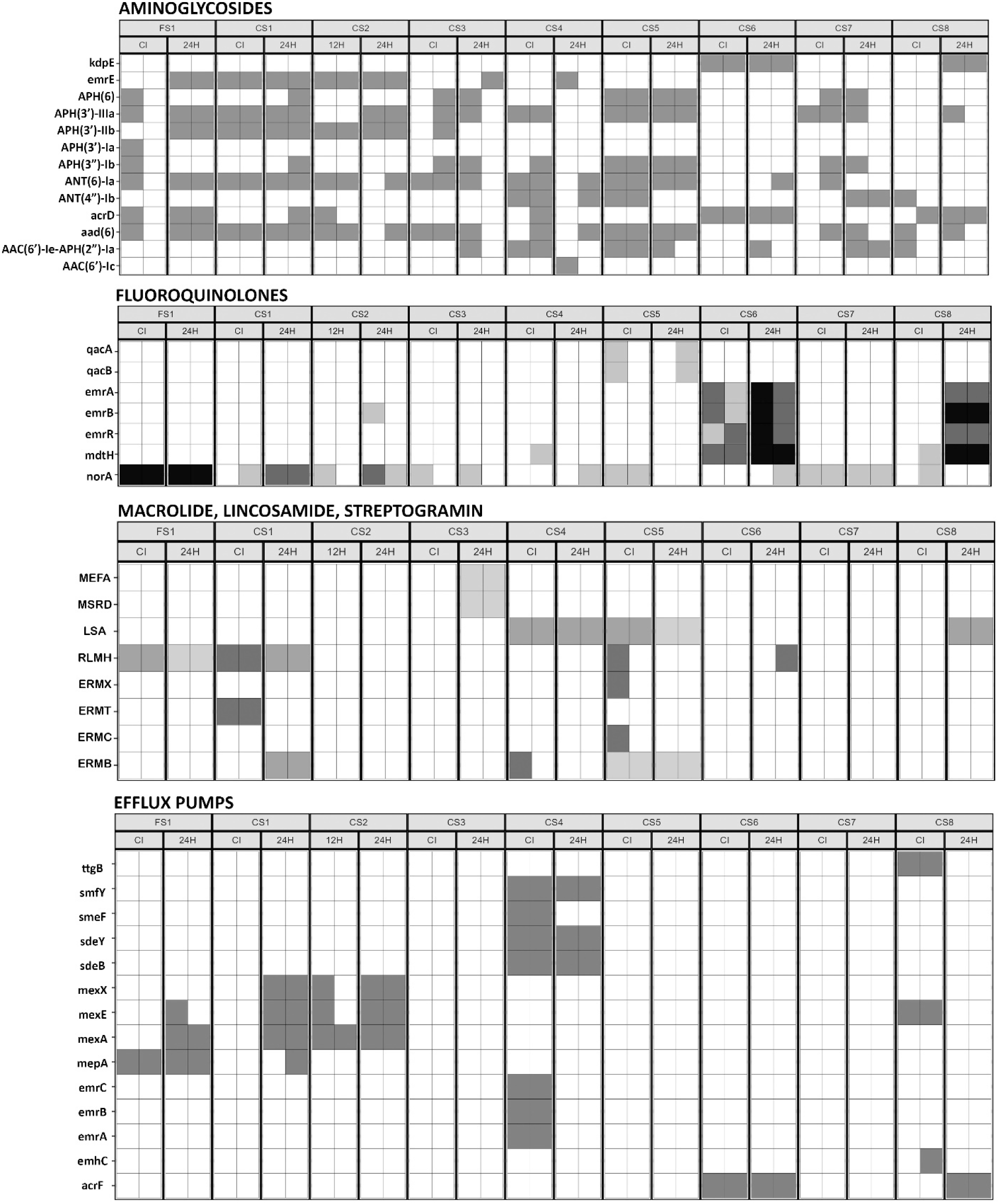
MGX Annotation of additional classes of AMR in feline and canine skin samples.

The National Antimicrobial Resistance Monitoring System (NARMS) of the Center for Veterinary Medicine at the FDA actively monitors AMR associated with humans and animals. Collaboration with the Veterinary Laboratory Investigation and Response Network (Vet-LIRN) monitors AMR associated with bacteria isolated from companion animals in a coordinated network across the United States and Canada. A subset of genes represent clinically important antimicrobial MDR, XDR and DTR monitoring targets (31).

**Figure 6.**
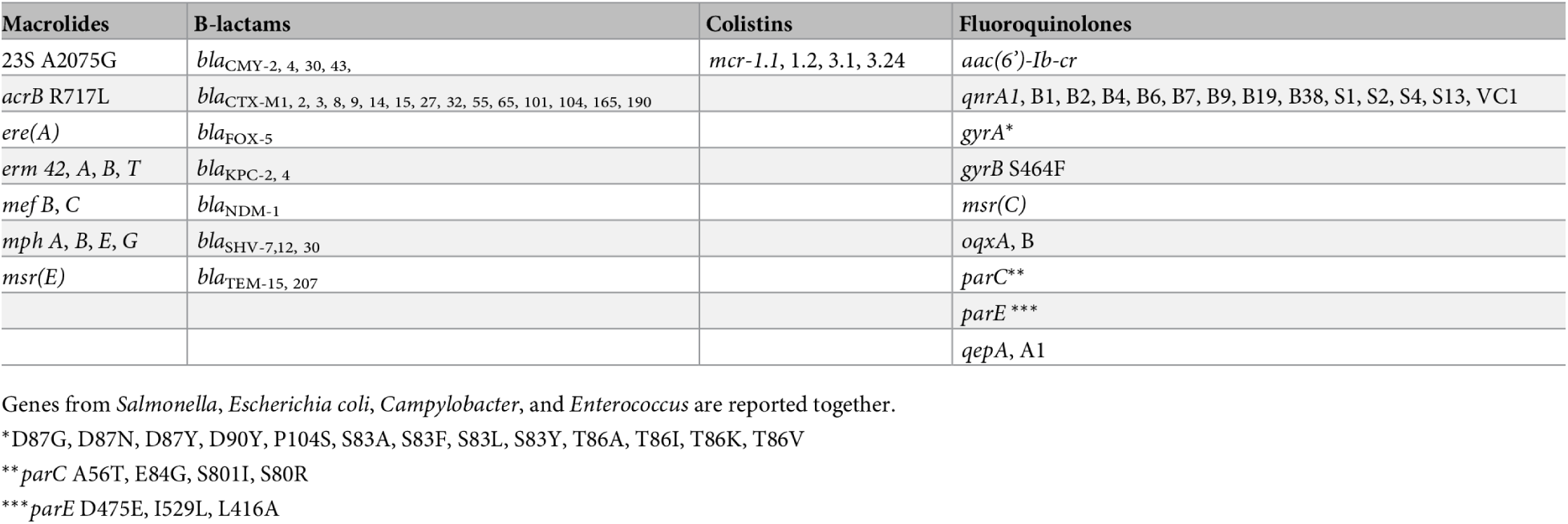
ARGs under active surveillance by NARMS.

In addition to the specific ARGS and pathogens monitored by NARMS (Figure 6), a larger list of pathogens is actively monitored by the European Antimicrobial Resistance Surveillance Network in Veterinary Medicine (EARS-Vet)(26). EARS-Vet is comprised of 38 partners spanning 18 countries and coordinates with the World Health Organization (WHO) to monitor 11 species known to cause disease in animals including *Escherichia coli, Klebsiella pneumoniae, Mannheimia haemolytica, Pasteurella multocida, Actinobacillus pleuropneumoniae, Staphylococcus aureus, Staphylococcus pseudintermedius, Staphylococcus hyicus, Streptococcus uberis, Streptococcus dysgalactiae, and Streptococcus suis.* An added benefit of metagenomic data is the ability to identify these species with demonstrated zoonotic import in conjunction with ARGs.

**Figure 7.**
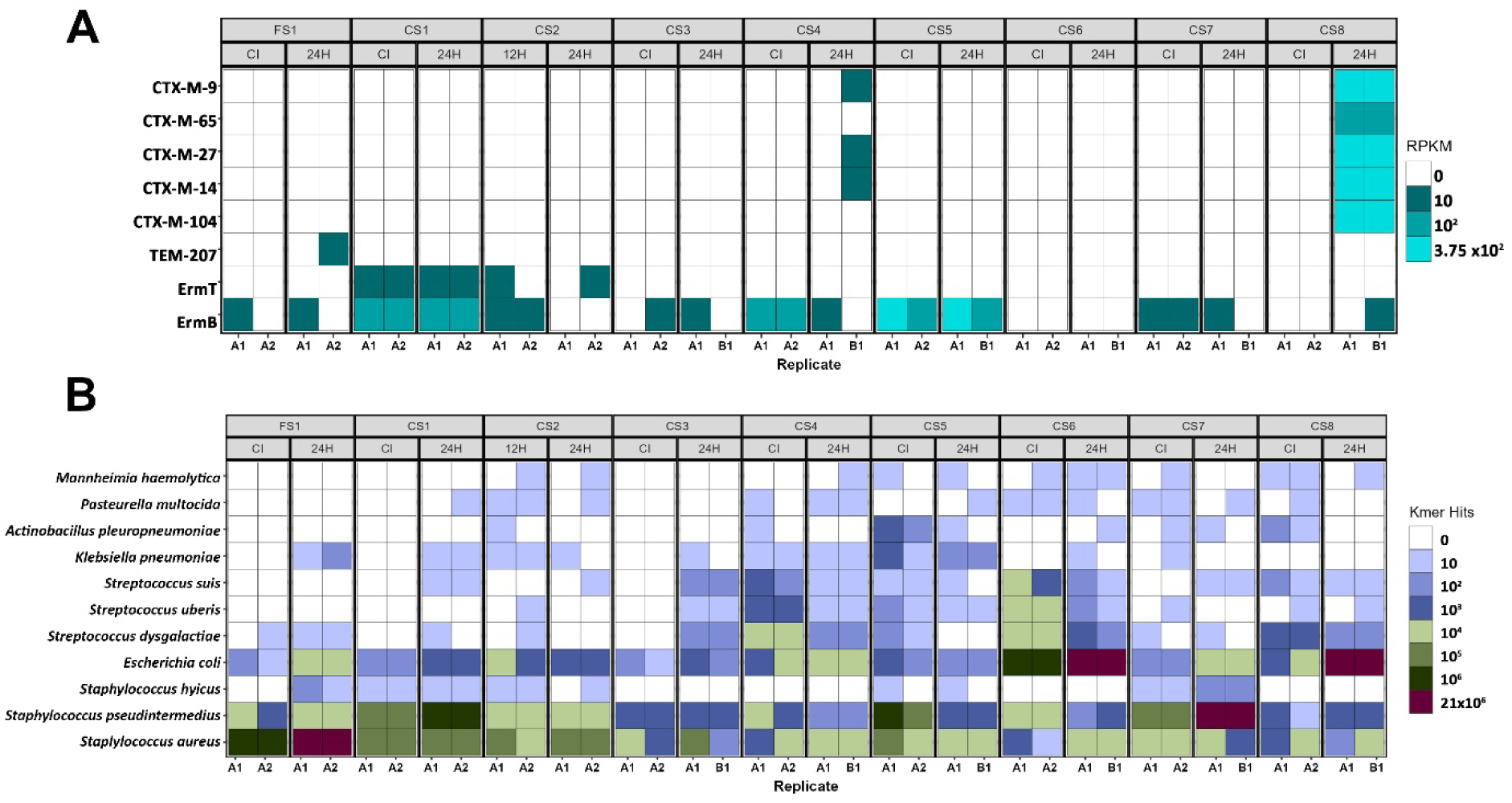
Surveillance utility of MGX data: NARMS ARG targets and EARS-Vet Pathogens observed in Feline and Canine skin samples.

### Plasmids

Plasmid data were available for five of the nine skin infection samples. The following replicon types were observed in feline and canine skin samples:

- **FS1**: IncI1, IncI2, IncP, IncA/C, IncA/C-like, IncFII(pKPX-1), Inc18
- **CS1**: pSK41-derivative (integrated)
- **CS4**: pSK41 (Gram-positive)
- **CS5**: RepUS11, RepUS14 (mobilizable)
- **CS8**: IncFII, IncFIB

The IncI, IncI2, IncA/C, IncF and IncP plasmid families are predominant Gram-negative AMR vectors across human and animal monitoring efforts. The pSK41-family plasmids are Gram-positive conjugative multi-resistance replicons in Staphylococci. Pheromone-responsive RepUS11 plasmids (e.g., cfr/cyl) are a growing concern because they connect linezolid resistance with high-level virulence in *Enterococcus.* Monitoring plasmids and gene carriage provides insight into transmissible resistance and selection factors. Full details of replicon families of identified plasmids with typical drug resistance classes and links to known strains of clinical importance are listed in Table 3.

**Table 3.** Replicon families of plasmids identified across the skin samples with characteristic ARG carriage (not necessarily observed in the samples evaluated here).

| Skin ID | Replicon Family | Drug Resistance Classes | Taxa Strain Association |
| --- | --- | --- | --- |
| FS1 | IncI2 | Polymyxins (mcr-1) | *E. coli*; IncI2 is dominant in mcr-1 global dissemination |
| FS1 | IncA/C-like | AmpC Beta-lactams (CMY-2), Phenicol, Sulfonamides, Tetracyclines | Salmonella Reading; 2019 U.S. outbreak in turkey products |
| FS1 | IncA/C | AmpC Beta-lactams (CMY-2), Phenicol, Sulfonamides, Tetracyclines | MDR Salmonella Newport; zoonotic foodborne outbreaks |
| FS1 | IncI1 | Tetracyclines, Streptomycin, Sulfonamides | Prototype IncI1 plasmid in Salmonella Typhimurium |
| FS1 | IncFII(pKPX-1) | Carbapenems (KPC-2) | *K. pneumoniae*; common in U.S. hospital outbreaks |
| FS1 | IncP | Polymyxins (mcr-3), Carbapenems (NDM-5) | *E. coli* ST206; rare but high-risk dual resistance |
| FS1 | Inc18 | Glycopeptides (vanA), Aminoglycosides, Macrolides | *Enterococcus faecium*; Inc18 vanA plasmid circulating in hospitals |
| CS1 | pSK41-derivative (integrated) | Trimethoprim, Tetracyclines, MLS<sub>B</sub> | Livestock-associated MRSA ST398, chromosomally integrated but mobile |
| CS4 | pSK41 (Gram-positive) | Aminoglycosides, MLS<sub>B</sub>, Tetracyclines, Cadmium | Common in MRSA; conjugative plasmid in *S. aureus* and CoNS |
| CS5 | RepUS11 | Oxazolidinones (linezolid), Cytolysin | *Enterococcus faecalis* OG1RF; virulence + resistance |
| CS5 | RepUS14 (mobilizable) | Tetracyclines | Found in *Staphylococcus aureus* SA01 |
| CS8 | IncFII | Extended-spectrum Beta-lactams (CTX-M) | *E. coli*; linked to community and hospital outbreaks |
| CS8 | IncFIB | Extended-spectrum Beta-lactams (CTX-M-15), Fluoroquinolone efflux (oqxAB) | *K. pneumoniae* ST383; high-risk clone |

**Table 4.** Plasmid-Associated Resistance Patterns.

| Plasmid Type | Samples | Primary Resistance Mechanisms |
| --- | --- | --- |
| IncFII/IncFIB | FS1, CS8 | optrA/poxTA (linezolid),<br>TMP/sulfa resistance |
| pSK41-related | CS1, CS4 | MLS_B, tetracycline,<br>aminoglycoside resistance |
| Multiple Inc types | FS1 | Broad-spectrum resistance<br>including tet, cat, arr genes |

Using data from VDL culture and AST results, clinical treatments were proposed by internal FDA AI (Initial Recommendation). With the addition of genes identified by MGX data, a new column of revised recommendations was returned which demonstrates one of the ways in which MGX data may be able to contribute to AI diagnostics in the future (Table 5).

**Table 5.**
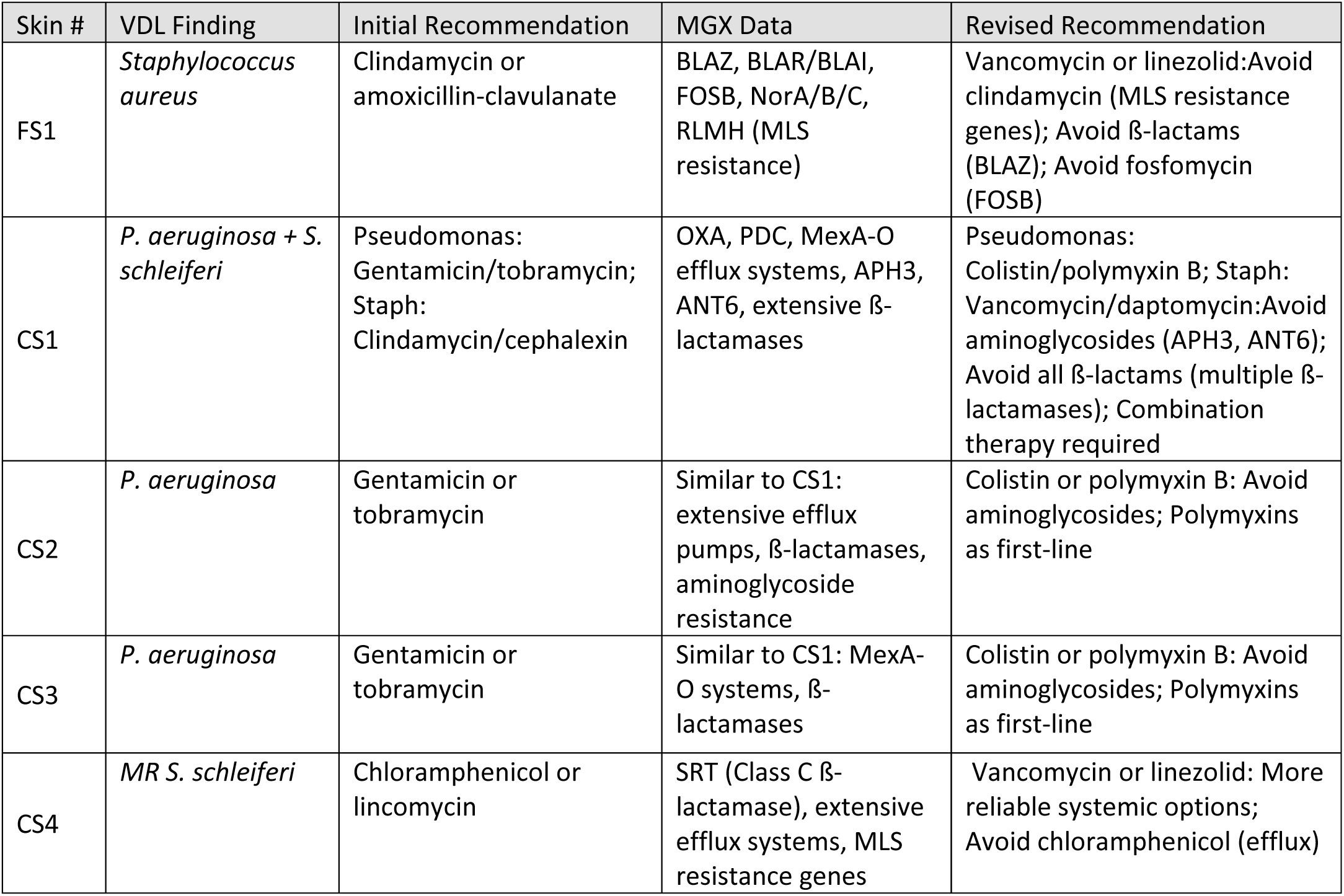
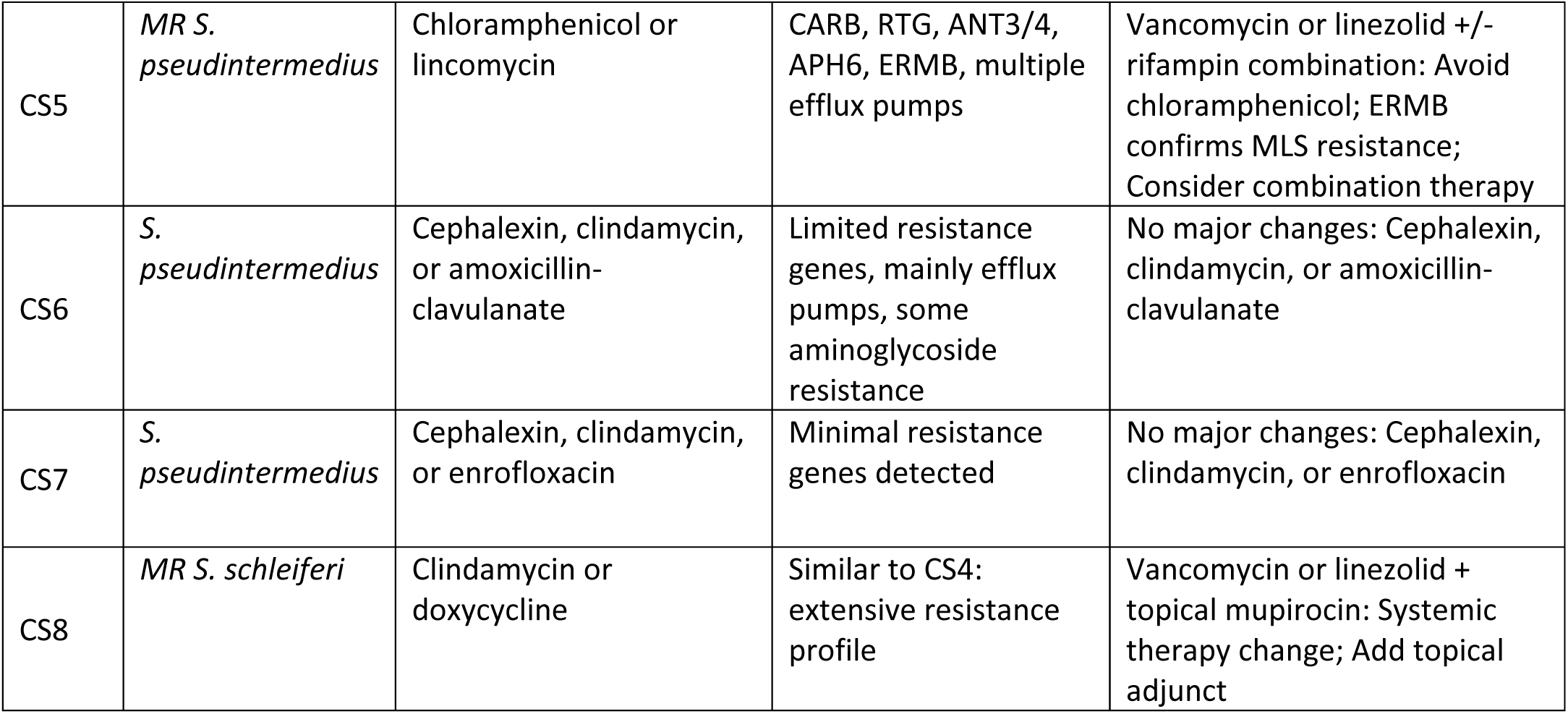
VDL and MGX results and how they may influence treatment.

| Skin # | VDL Finding | Initial Recommendation | MGX Data | Revised Recommendation |
| --- | --- | --- | --- | --- |
| FS1 | *Staphylococcus aureus* | Clindamycin or amoxicillin-clavulanate | BLAZ, BLAR/BLAI, FOSB, NorA/B/C, RLMH (MLS resistance) | Vancomycin or linezolid: Avoid clindamycin (MLS resistance genes); Avoid β-lactams (BLAZ); Avoid fosfomycin (FOSB) |
| CS1 | *P. aeruginosa* + *S. schleiferi* | Pseudomonas: Gentamicin/tobramycin; Staph: Clindamycin/cephalexin | OXA, PDC, MexA-O efflux systems, APH3, ANT6, extensive β-lactamases | Pseudomonas: Colistin/polymyxin B; Staph: Vancomycin/daptomycin: Avoid aminoglycosides (APH3, ANT6); Avoid all β-lactams (multiple β-lactamases); Combination therapy required |
| CS2 | *P. aeruginosa* | Gentamicin or tobramycin | Similar to CS1: extensive efflux pumps, β-lactamases, aminoglycoside resistance | Colistin or polymyxin B: Avoid aminoglycosides; Polymyxins as first-line |
| CS3 | *P. aeruginosa* | Gentamicin or tobramycin | Similar to CS1: MexA-O systems, β-lactamases | Colistin or polymyxin B: Avoid aminoglycosides; Polymyxins as first-line |
| CS4 | *MR S. schleiferi* | Chloramphenicol or lincomycin | SRT (Class C β-lactamase), extensive efflux systems, MLS resistance genes | Vancomycin or linezolid: More reliable systemic options; Avoid chloramphenicol (efflux) |
| CS5 | *MR S. pseudintermedius* | Chloramphenicol or lincomycin | CARB, RTG, ANT3/4, APH6, ERMB, multiple efflux pumps | Vancomycin or linezolid +/- rifampin combination: Avoid chloramphenicol; ERMB confirms MLS resistance; Consider combination therapy |
| CS6 | *S. pseudintermedius* | Cephalexin, clindamycin, or amoxicillin-clavulanate | Limited resistance genes, mainly efflux pumps, some aminoglycoside resistance | No major changes: Cephalexin, clindamycin, or amoxicillin-clavulanate |
| CS7 | *S. pseudintermedius* | Cephalexin, clindamycin, or enrofloxacin | Minimal resistance genes detected | No major changes: Cephalexin, clindamycin, or enrofloxacin |
| CS8 | *MR S. schleiferi* | Clindamycin or doxycycline | Similar to CS4: extensive resistance profile | Vancomycin or linezolid + topical mupirocin: Systemic therapy change; Add topical adjunct |

### Functional gene data

The accessibility of biomarkers for diagnostic etiologies, features, or disease states based on functional metabolic genes is rapidly changing the way in which patients are evaluated and managed for certain conditions. If specific biomarkers could be linked to *Staphylococcus* infections to distinguish them from *Pseudomonas* infections, that could potentially expedite initial treatment. We observed only one significant differentially abundant metabolic feature between *Staphylococcus* and *Pseudomonas* infections with 4-deoxy-l’threo-hex-4-enopyranuronate (with the caveat that sample size in this case study does not necessarily support biological relevance). The differential incidence of genes for production of uronic acid (4-deoxy-l’threo-hex-4-enopyranuronate) were significantly elevated in the *Staphylococcal* infections contrasted to the *Pseudomonas* infections (Figure 8).

**Figure 8.**
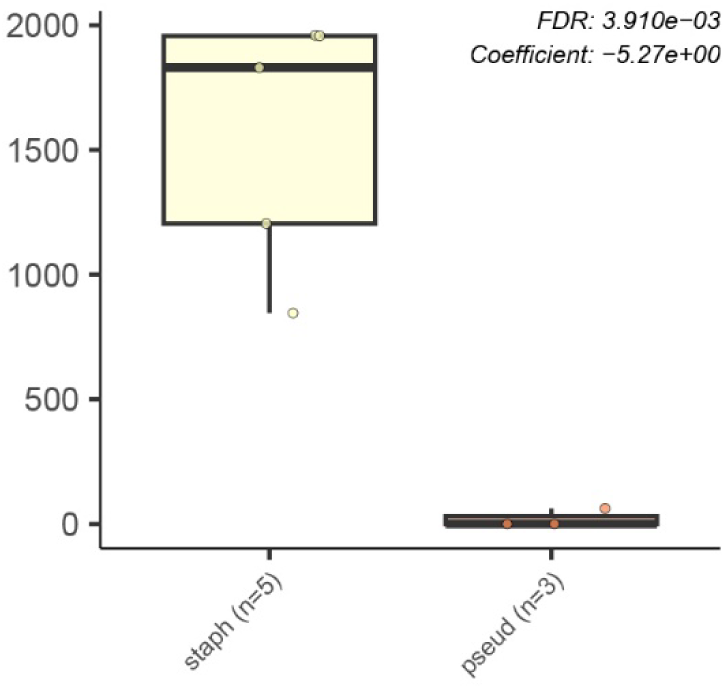
4-deoxy-l’threo-hex-4-enopyranuronate is differentially abundant in *Staphylococcal* infections.

### AI predictions

Metagenomic data provides an enormous, at times overwhelming amount of data describing taxonomy, AMR, and metabolomic features of samples. The work presented here focuses on how these data may be integrated into assays to advance veterinary diagnostics. To this end, we also explored several models to evaluate how MGX data may predict antimicrobial susceptibility or resistance. To predict methicillin resistance, the top 32 species (excluding MR species MRSS and MRSP) were used as input data. Data were split into training and test sets. 70% of the data was used to train the models and then the other 30% was tested to see if the model would correctly predict on that held out data. Results of the test data set showed perfect prediction, but this was almost certainly due to the very small test set and would likely not hold up using a larger set of data. A decision tree model to identify cutoff abundances in one or more species that may differentiate the samples produced an over simplified tree with only one split: classifying all samples with > 0.013 abundance of *Pantoea agglomerans* as having MR strains. This is again oversimplified due to the small dataset and may not have biological relevance. Only one sample did not follow the prediction pattern by *Pantoea agglomerans* abundance which was the feline sample, which actually supports the potential utility of the model. If training data was predominantly based on canine samples, it makes sense that the microbiome of the feline sample may not align with predominantly canine-microbiome based prediction.

## Discussion

In this case study, we evaluated metagenomic data in conjunction with state-of-the-art VDL aerobic culture and AST results for a small number of feline and canine skin samples. The results help define the immense array of opportunities for advancing One Health approaches to veterinary diagnostics. While too much data, without appropriate standards for quality and interpretation may actually confound rather than improve treatment (27), there exists an undeniable frontier of utility for MXG data in veterinary diagnostics.

An exciting application for the type of data presented here centers around provision of simultaneous pathogen, AMR, and plasmid identification which may, when appropriately optimized and validated, direct more precise and judicious antibiotic administration.

Another exciting opportunity provided by MGX data is the ability to identify cross domain linkage, co-occurrence, and biological relationships that may improve our understanding of currently uncharacterized infection ecologies. Information about co-occurring taxa and cross domain relationships has been a success story for treatment of heartworm (*Dirofilaria immitis*) in animals for decades (treating the bacterial symbiont *Wolbachia* with antimicrobial (doxycycline)) in conjunction with direct management of filaria (28, 29)) and for Lymphatic Filariasis where elimination of the bacterial symbiont (*Wolbachia*) of eukaryotic parasite *Wucheria bancrofti* served as an effective treatment for parasitic infections of humans(30–32). A dynamic and at times antagonistic interplay between *Malassezia* and *Staphylococcu*s has been described, including reduced susceptibility of *Malassezia* to the azole antifungal ketoconazole when co-cultured with *Staphylococcus*(33, 34). Co-occurrence and enrichment of both taxa has been observed in scalp microbiomes of human patients with seborrheic dermatitis(35) and *Malessezia* has been shown to antagonize *Staphylococcus* and disrupt its biofilm formation(36). A symbiotic relationship between *Malassezia pachydermatis* and commensal staphylococci has even been proposed(37). We observed both *Malassezia* and *Staphylococcus* in two infections. Perhaps intentional manipulation of *Malassezia* could impact survival of *Staphylococcus* in MR infections, providing new approaches when antibacterials are not effective.

MGX data may also be useful for improved characterization of prevalence and co-occurrence of zoonotic pathogens. Methicillin-resistant staphylococci (MRSA, MRSP, MRSS) can transfer between humans and pets making co-occurrence of these species in feline and canine infections important to understand. Additionally *Entamoeba dispar*(38) was observed in one canine samples and has reported ability to infect both humans and animals and virus RD1114 was also observed, which unlike typical endogenous retroviruses that are transmitted vertically through germline integration, has demonstrated capacity for horizontal transmission between species. Additionally, we highlight the potential for MGX data to support metabolomic or chemical biomarker discovery. Metabolomic data have differentiated between feline chronic enteropathy and small-cell lymphoma(39), and provided biomarkers for canine sepsis (40), sow pregnancy (41), and hyperketonaemia in dairy cows, to list a few. Metabolomic screens are expensive and complicated and often DNA data can identify differential incidence of pathways for key metabolites typically measured by chemical techniques(42, 43).

### Accelerated Cures

A primary objective for FDA is the acceleration of cures (44) for the American public. For the Center for Veterinary Medicine (CVM), this includes companion animals. Research at the complex One Health nexus of human, animal, and environmental health continually evaluates new alternative methods to advance human and animal health. Cures and treatment cannot be expedited if diagnostics do not sufficiently inform medical response. Even the small case study presented here highlights the vast frontier of opportunity for advancing veterinary diagnostics using MGX data. Genetic, taxonomic, and metabolomic features of infections may provide novel biomarkers to advance monitoring efforts for a broad range of conditions. Simultaneous identification of pathogen targets, co-occurring species, ARGs, plasmids, and metabolic and ecologic features will undoubtedly improve our ability to identify and steward antimicrobial resistance transmission. Additionally, MGX data input to AI models shows exciting promise for advancing diagnostics and identifying significant linkages. MGX data, collected in coordination with VDL pathogen assays and carefully curated metadata, demonstrates the potential to advance our ability to predict both methicillin resistance, pathogen source attribution, and features which may distinguish between infection types. These data have the potential to inform epidemiology, identify novel biomarkers for improved surveillance and risk assessment, as well as advance precision treatment strategies.

## Materials and Methods

### DNA Extraction

Skin was swabbed by a veterinarian with Amies non charcoal swabs and stored at -20°at UPenn Veterinary Hospital and shipped to FDA Center for Veterinary Medicine labs for pilot MGX analyses. Swabs were cut in half and DNA was extracted from half which received no enrichment and half after 24H enrichment in universal pre-enrichment broth (UPB) at 37° using the Qiagen DNeasy Blood and Tissue Kit according to the manufacturer’s specifications.

### Library Preparation and Sequencing

DNA libraries was prepared using the Illumina DNA Library Prep Kit according to the manufacturers specifications (Illumina).

https://www.protocols.io/edit/illumina-dna-prep-sop-bzstp6en

Sequencing was performed on a NextSeq 2000 with 2 x 150 cycles using the NextSeq 1000/2000 P2 Reagent Kit (300 Cycles). Libraries were diluted to a 750 pM loading concentration according to the Illumina’s specifications (ext-link ext-link-type="uri" xlink:href="https://support.illumina.com/content/dam/illumina-support/documents/documentation/system_documentation/NextSeq2000/nextseq-1000-2000-sequencing-system-guide-1000000109376-05.pdf">NextSeq Denature and Dilute Libraries Guide).

### Bioinformatic Analysis

Sequencing data was demultiplexed (bcl2fastq2), screened/trimmed using Trimmomatic (45). Quality-checked reads resulted in an average of 22 million reads per sample for further downstream analyses.

### AMR Annotation

Paired end FASTQ files were analyzed using the AMR++ pipeline(46) with the MEGARes database v2 using default parameters. Additionally, the AMRFinder Plus database was used for AMR annotation using SAUTE (47) on the FDA Human Foods Program (HFP) high performance cluster(HPC). https://github.com/ncbi/amr/wiki/Methods and BLAST was used with the CARD(48) database to annotate sequence data according to default parameters. Reads were also evaluated using the COSMOS ID analytical pipeline (AMR database update May 2025 https://www.cosmosid.com) to take contrast with in-house results). Counts and abundances from AMR annotation outputs were ‘normalized’ using scripts to assess ‘reads per kilobase of transcript’ (RPKM) to normalize gene reporting between different sites by accommodating for variation in number of sequencing reads per sample and gene length. Total reads in the sample were divided by 1,000,000 “per million” scaling factor to normalize for sequencing depth and provide ‘reads per million’ (RPM). RPM values were then divided by length of each gene in kilobases to report ‘RPKM.

https://github.com/SethCommichaux/AMRplusplus

Identification of plasmids was accomplished using Platon (49) with BLAST NCBI tools(50) to top hits.

### Bacterial Annotation

Determination of bacterial composition from shotgun sequencing was conducted using custom C++ programs developed to compile a *k*-mer signature database containing multiple unique 30 bp sequences per species and then identify each read in the input file using the 30 bp probes. The *k*-mer database used for this work contains 5900 target taxonomic entries, each consisting of approximately 40,000 (range 44 to 80,000) unique *k*-mers. The database includes 1100 different bacterial genera, and 3500 species. Normalization is performed to correct for bias due to differing number of *k*-mers used per database entry and results are tabulated as percent of identified reads for each database entry. Kraken(51) is also used as a first pass taxonomic classifier.

### Fungal, Virus and Phage Annotation

The same method described above was created to annotate fungal and viral taxa from sequencing reads. Reads from annotations were double-checked by Blasting them against genomes of identified organisms to better assess the quality of the annotations. Additionally, the COSMOS ID analytical pipeline with the fungal database was used to contrast with results from the FDA in-house fungal annotation pipeline and database (https://www.cosmosid.com).

### Data Reporting and Visualization

Pipeline annotation outputs were visualized using R Studio (version 1.4.1717, using R version 4.1.1). Visualizations were created using ggplot2(52) and Adobe Photoshop.

## List of Abbreviations

AMR: Antimicrobial Resistance
ARG: Antimicrobial Resistant Gene
AST: Antimicrobial Susceptibility Testing
DTR: Difficult to Treat Resistance
EARS: Vet – European Antimicrobial
FDA: Food and Drug Administration
HGT: Horizontal Gene Transfer
MGX: Metagenomic
MDR: Multi-Drug Resistant
MRSP: Methicillin resistant *Staphylococcus pseudintermedius*
MR: Methicillin Resistant
MRSS: Methicillin Resistant *Staphylococcus schleiferi*
NARMS: National Antimicrobial Resistance Monitoring System
NGS: Next Generation Sequencing
PCR: Polymerase Chain Reaction
qMGX: quasiMGXs
VDL: Veterinary Diagnostic laboratories
WHO: World Health Organization

## Availability of Data

All sequences have been deposited in CVM Metagenomes in BioProject: PRJNA1335144 https://www.ncbi.nlm.nih.gov/bioproject/PRJNA1335144

## Authors’ contributions

AO, OC, SC, and SR conceived of the study. JD performed microbiological lab work. AO, SS, and BK conducted molecular lab work. MM, BK, AO, performed bioinformatic and statistical analyses. BK and AO created scientific visualizations. SC, SR, SP, and OC provided editorial guidance.

## Footnotes

1 ‘DTR’ is a newly recommended descriptor by the Global Antibiotic Research and Development Partnership (GARDP) to replace ‘MDR’ for better alignment with clinical decisions

